# Synthesis and validation of click-modified of NOD1/2 agonists

**DOI:** 10.1101/2023.03.28.534546

**Authors:** Ravi Bharadwaj, Madison V. Anonick, Siavash Mashayekh, Ashley Brown, Kimberly A. Wodzanowski, Kendi Okuda, Neal Silverman, Catherine L. Grimes

## Abstract

NOD1 and NOD2 sense small bacterial peptidoglycan fragments often called muropeptides. These muropeptides include iE-DAP and MDP, the minimal agonists for NOD1 and NOD2, respectively. Here, we synthesized and validated alkyne-modified muropeptides, iE-DAP-Alk and MDP-Alk, for use in click-chemistry reactions. While it has long been known that many cell types respond to extracellular exposure to muropeptides, it is unclear how these innate immune activators access their cytosolic innate immune receptors, NOD1 and NOD2. The subcellular trafficking and transport mechanisms by which muropeptides access these cytosolic innate immune receptors are a major gap in our understanding of these critical host responses. The clickchemistry-enabled agonists developed here will be particularly powerful to decipher the underlying cell biology and biochemistry of NOD1 and NOD2 innate immune sensing.

## Introduction

Innate immune cells such as macrophages, neutrophils, NK cells, and dendritic cells harbor specific innate immune receptors present on the cell surface, within intracellular compartments, or in the cytosol. Toll-like receptors (TLR) were the earliest discovered and most extensively studied cell surface and vesicular innate immune receptors [1]. However, cytosolic innate immune receptors play an equally critical role in host defense by sensing and discriminating between pathogenic and commensal microbes [1]. This category of receptors includes several proteins capable of sensing nucleic acids, such as cGAS, RIG-I, NLRP1, and AIM2, recognizing bacterial products (NOD1/2, NAIPs/NLRC4) or other danger signals (NLRP1/3)[2–5]. Among these cytosolic receptors, NOD1 and NOD2 sense small bacterial cell wall fragments, often referred to as muropeptides, that access the intracellular environment [6].

Peptidoglycan (PGN), the major constituent of the bacterial cell wall, is a network of long carbohydrate chains cross-linked via short-stem peptides, that provides the bacterium with mechanical strength and protection from internal and external stressors[7]. Muropeptides are often shed into the environment during cell wall turnover and biosynthesis, and trigger NOD1 or NOD2 upon gaining access to the cytosol [8]. For example, muramyl dipeptide (MDP) a major component of Freund’s adjuvant, is the minimal molecular motif that activates NOD2 triggering an NF-κB inflammatory response[9], although recent studies have shown that MDP must be first phosphorylated, by the host cytosolic enzyme *N*-acetylglucosamine kinase (NAGK), in order to activate NOD2 [10]. Other muropeptides that contain *meso*-diaminopimelic acid (*m*-DAP), an amino acid unique to Gram-negative bacteria and Gram-positive bacilli peptidoglycan, activate NOD1 [11]. Recently, it has shown that mice lacking NOD1/2 receptors are more susceptible to intracellular and extracellular bacterial infections including *H. pylori, Clostridium difficile, Listeria monocytogenes, Chlamydia pneumoniae*, and *Staphylococcus aureus* [12–20]. NOD2 is also strongly linked to inflammatory bowel disease [21] and several studies found that NOD2 is critical for the healthy gut epithelial structure and immunity [6]. Many gaps exist in our current knowledge of NOD1/2 pathways, such as if phosphorylation of all muropeptides is required for their NOD1/2 agonist activity, how different muropeptides access the cytosol to trigger NOD1 or NOD2, what other proteins are involved in binding and sensing these muropeptides, and if different agonists trigger distinct signaling pathways.

Click chemistry is a powerful tool in the field of drug discovery, cell signaling, and pharmaceutical sciences[22]. This tool provides a rapid means to synthesize biomolecules useful in high throughput screening, examining biomolecular interactions, and fluorescent microscopic detection [23]. While several types of click reactions have been developed, Cu^I^-catalyzed Huisgen 1,3-dipolar cycloaddition of azides and terminal alkynes is the most widely implemented in the synthesis of natural products with increased efficiency and biologically compatible reagents [24, 25]. Using this approach, we have generated “click” modified D-isoGlutamate-*meso*-Diaminopimelic acid alkyne (iE-DAP-Alk) and 2-alkyne muramyl dipeptide (MDP-Alk), which represent the minimal NOD1 and NOD2 agonists respectively [26]. We note that others have made other PGN-related probes. Howard Hang and co-workers used a similar approach in their synthesis of chemo-proteomic PGN probes [27], combining photoactivatable crosslinkers with a clickable alkyne moiety; while others have used solid phase synthesis to incorporate click probes on PGNs [28]. A recent review from Tanner and co-workers succinctly highlights our and others’ synthetic efforts in this space [29].

Here, we report the chemical synthesis of iE-DAP-Alk and MDP-Alk, for use in clickchemistry reactions. This modification is smaller than those previously reported, and valuable in recent work where we probed the biological role of PGN transporters [30]. The modified iE-DAP and MDP showed similar potency as the native molecules in the NF-κB reporter assays in transfected cell lines, and in cytokine induction in IFNγ-primed bone marrow-derived macrophages (BMDM). Using these click-muropeptides and fluorescent confocal microscopy we have also confirmed the internalization of iE-DAP and MDP in WT BMDM and HCT116 cells.

## Methods

### Synthesis of iE-DAP-Alk

Synthesis of iE-DAP-Alk follows an eight-step process (all of which have literature precedent [27,28, 37–39, 44]). Overall chemical synthesis of iE-DAP-Alk is described below and presented in **Figure 1**. We note that the synthesis of intermediate 5 follows prior literature while transformation from 5 to 8 is supported by other related literature for the synthesis of similar molecules. The synthesis is detailed below:

**Figure 1.**
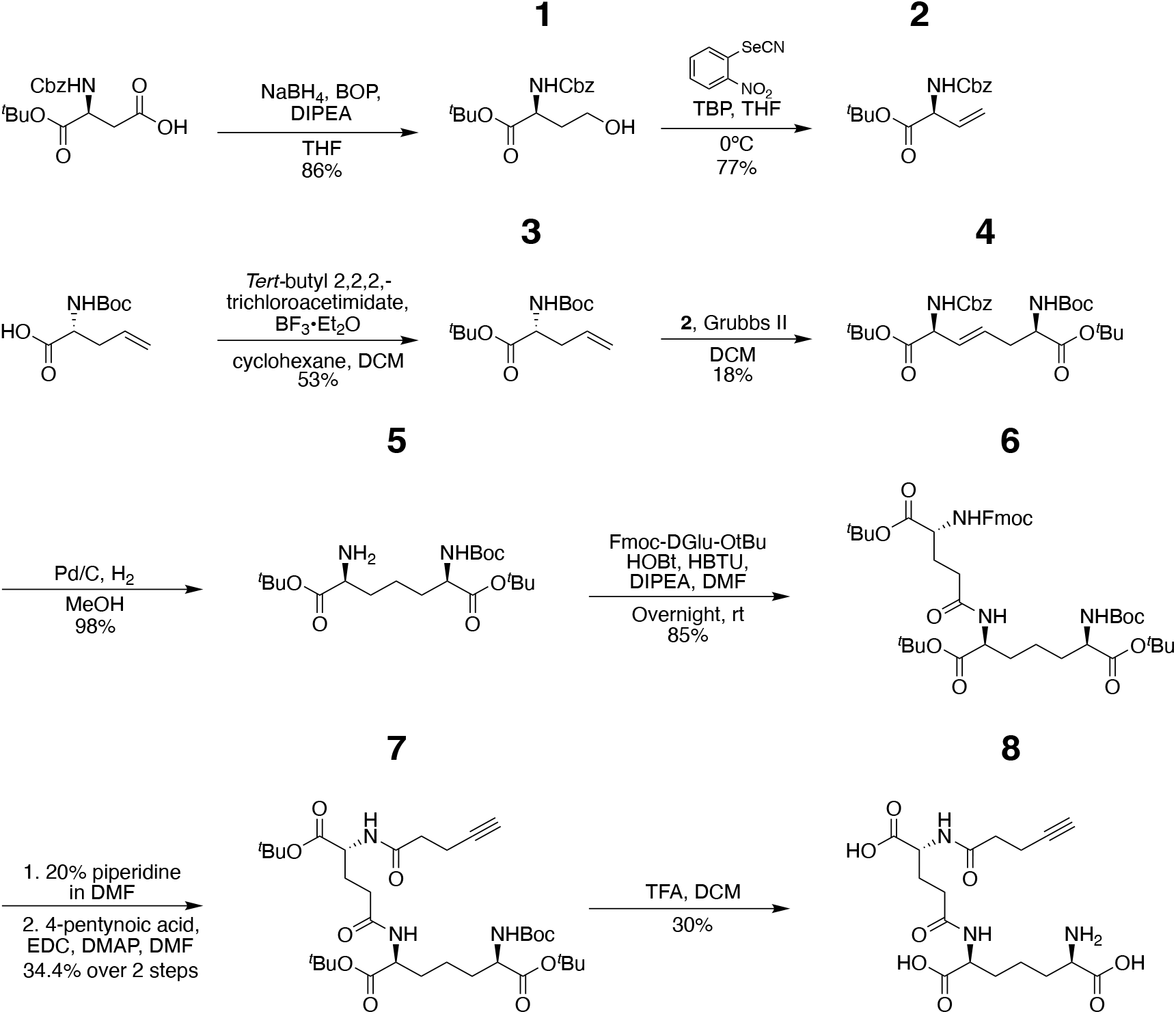
Schematic overview of synthesis of iE-DAP-Alk. The overall strategy for chemical synthesis of iE-DAP-alkyne with biologically active stereochemistry followed literature precedent; the protocols presented in this manuscript give critical details for this specific workflow, making it easier for the production of these important PGN probes.

#### *Tert-butyl* ((benzyloxy)carbonyl)-*L*-homoserinate (1)

To a suspension of Z-l-Asp-a-OtBu (2.04 g, 6.32 mmol, 1.0 eq.) and benzotriazol-1-yl-oxy-tris(dimethylamino)phosphonium hexafluorophosphatein (BOP) (3.47 g, 7.84 mmol, 1.2 eq.) in anhydrous THF (50 ml) was added DIPEA (1.43 ml, 8.2 mmol, 1.3 eq.). The mixture stirred for 10 min at room temperature before cooling down the reaction to 0°C. Then, sodium borohydride (NaBH_4_) (312 mg, 8.2 mmol, 1.3 eq.) was added in two portions. The reaction stirred overnight at room temperature. The reaction was then condensed, diluted with ethyl acetate and washed with 1N HCl (thrice), sat. sodium bicarbonate (thrice), brine (once) and dried over sodium sulfate. After concentration, the crude product was purified using flash chromatography (hexane/ethyl acetate-gradient from 5% to 20% ethyl acetate) to give **1** as a colorless oil, matching those previously reported in the literature [31]. (1.55 g, 86%). ^1^H NMR (600 MHz, MeOD) δ 7.41 – 7.23 (m, 5H), 5.13 – 5.02 (m, 2H), 4.20 (dd, *J* = 9.7, 4.7 Hz, 1H), 3.69 – 3.57 (m, 2H), 2.06 – 1.97 (m, 1H), 1.85 – 1.77 (m, 1H), 1.45 (s, 7H), 1.38 (s, 1H). ^13^C NMR (151 MHz, MeOD) δ 173.52, 158.67, 138.25, 129.45, 128.99, 128.84, 82.70, 67.91, 67.58, 59.21, 53.59, 53.49, 35.24, 28.21. LRMS (ESI) m/z: [M + Na]^+^ Calcd for C_16_H_23_NNaO_5_ 332.14; Found 332.28. NMR spectra shown in Figure S1A

#### *Tert-butyl* (*S*)-2-(((benzyloxy)carbonyl)amino)but-3-enoate (2)

a solution of **1** (2.0 g, 6.46 mmol, 1.0 eq.) in anhydrous THF (32 mL) was cooled down to 0°C in an ice bath. Tri-n-butylphosphine (TBP) (1.8 mL, 7.11 mmol, 1.1 eq.) was added dropwise followed by 2-nitrophenyl selenocyanate (1.61 g, 7.11 mmol, 1.1 eq.). The reaction was warmed to room temperature and stirred for 1 h. The reaction was cooled down to 0°C again and 30% hydrogen peroxide solution (2.53 mL) was added. The reaction stirred overnight at room temperature. The reaction was then poured into saturated sodium chloride (60 ml) and extracted with ethyl acetate (thrice). Combined organics were washed with 1N HCl (thrice), washed with brine (once) and dried over sodium sulfate. After concentration, the crude product was purified using flash chromatography (hexane/ethyl acetate gradient from 0% to 30% ethyl acetate) to give **2** as a yellow oil, matching those previously reported in the literature [32]. (1.45 g, 77%). ^1^H NMR (400 MHz, MeOD) δ 7.41 – 7.25 (m, 6H), 5.93 (ddd, *J* = 16.7, 10.4, 5.9 Hz, 1H), 5.39 – 5.31 (m, 1H), 5.28 – 5.21 (m, 1H), 5.10 (t, *J* = 2.2 Hz, 2H), 4.69 – 4.62 (m, 1H), 1.45 (s, 7H), 1.38 (s, 1H). ^13^C NMR (101 MHz, MeOD) δ 171.35, 158.32, 138.19, 133.83, 129.46, 129.02, 128.86, 118.01, 83.13, 67.69, 58.72, 28.16. LRMS (ESI) m/z: [M + Na]^+^ Calcd for C_16_H_21_NNaO_4_ 314.13; Found 314.43. NMR spectra shown in Figure S1B.

#### *Tert*-butyl (*R*)-2-((*tert*-butoxycarbonyl)amino)pent-4-enoate (3)

To a solution of d - 2-allyl-glycine-α-tertbutyl (0.932 g, 4.33 mmol, 1.0 eq.) in anhydrous DCM (4.33 mL) was added the solution of *tert*-butyl 2,2,2-trichloroacetimidate (1.9 g, 8.66 mmol, 2.0 eq.) in anhydrous cyclohexane (28 mL) dropwise followed by addition of boron trifluoride diethyl etherate (100 μL). Upon reaction completion, solid NaHCO_3_ was added; the reaction was then filtered, and the solvent was evaporated. The crude product was then purified using flash chromatography (hexane/ethyl acetate-gradient from 0% to 15% ethyl acetate) to give **3** (617 mg, 53%), matching those previously reported in the literature [33]. ^1^H NMR (400 MHz, MeOD) δ 5.85 – 5.70 (m, 1H), 5.17 – 5.04 (m, 3H), 4.03 (dd, *J* = 7.9, 5.5 Hz, 1H), 2.53 – 2.45 (m, 1H), 2.43 – 2.34 (m, 1H), 1.46 (s, 9H), 1.44 (s, 11H). ^13^C NMR (101 MHz, MeOD) δ 172.91, 157.84, 134.61, 118.51, 82.67, 80.46, 56.50, 55.43, 37.20, 28.71, 28.29. LRMS (ESI) m/z: [M + H]^+^ Calcd for C_14_H_26_NO_4_ 272.18; Found 272.20. NMR spectra shown in Figure S1C

#### Di-*tert*-butyl(2*S*,6*R*,*E*)-2-(((benzyloxy)carbonyl)amino)-6-((*tert*-butoxycarbonyl) amino)hept-3-enedioate (4)

**2 (**458 mg, 1.57 mmol. 1.8 eq.) and **3** (237 mg, 0.87 mmol. 1.0 eq.) were dissolved in degassed anhydrous DCM (16 mL) and cannula transferred to an oven-dried round bottom flask charged with Grubbs ’catalyst 2^nd^ generation (66.6 mg, 0.078 mmol. 0.05 eq) under nitrogen. The reaction stirred 48 hours monitored by MS. The reaction was then concentrated, and the crude product was purified using flash chromatography (hexane/ethyl acetate 0.01% Triethylamine - gradient from 0% to 12% ethyl acetate) to give **4** as a yellow oil (148 mg, 18%)[34], matching those previously reported in the literature [34]. ^1^H NMR (400 MHz, MeOD) δ 7.40 – 7.25 (m, 5H), 5.80 – 5.70 (m, 1H), 5.69 – 5.60 (m, 1H), 5.10 (s, 2H), 4.60 (d, *J* = 6.0 Hz, 1H), 4.04 (dd, *J* = 7.6, 5.3 Hz, 1H), 2.53 – 2.37 (m, 2H), 1.45 (s, 14H), 1.43 (s, 10H). ^13^C NMR (101 MHz, MeOD) δ 172.66, 171.51, 158.22, 157.84, 138.16, 130.05, 129.46, 129.02, 128.88, 83.15, 82.85, 80.61, 67.69, 57.94, 55.33, 49.85, 49.64, 49.43, 49.21, 49.00, 48.79, 48.57, 48.36, 35.50, 28.73, 28.29, 28.23.LRMS (ESI) m/z: [M + Na]^+^ Calcd for C_28_H_42_N_2_NaO_8_ 557.28; Found 557.23. NMR spectra shown in Figure S1D

#### Di-*tert*-butyl(2*S*,6*R*)-2-amino-6-((*tert*-butoxycarbonyl)amino)heptanedioate (5)

**4** (453 mg, 0.84 mmol. 1.0 eq.) was dissolved in methanol (25 mL) and Pd/C catalyst (100 mg, 10% loading) was added. The reaction stirred under H_2_ (balloon) overnight. The catalyst was then filtered over a pad of Celite, and the filtrate was condensed to give **5** as a colorless oil (330 mg, 98%).^1^H NMR (600 MHz, MeOD) δ 3.94 (dd, *J* = 9.0, 5.2 Hz, 1H), 2.96 (dd, *J* = 9.0, 5.5 Hz, 1H), 2.33 (s, 6H), 1.80 – 1.67 (m, 2H), 1.67 – 1.56 (m, 2H), 1.49 (s, 9H), 1.46 (s, 7H), 1.44 (s, 11H). ^13^C NMR (151 MHz, MeOD) δ 173.70, 172.85, 82.58, 82.53, 80.42, 69.73, 55.62, 49.43, 49.28, 49.14, 49.00, 48.86, 48.72, 48.57, 42.09, 32.48, 30.40, 28.75, 28.52, 28.39, 28.27, 23.52. LRMS (ESI) m/z: [M + H]^+^ Calcd for C_20_H_39_N_2_O_6_ 403.28; Found 403.27. NMR spectra shown in Figure S1E

#### Tri-*tert*-butyl (5*R*,10*S*,14*R*)-1-(9*H*-fluoren-9-ylmethoxycarbonyl)-18,18 dimethyl-3,8,16-trioxo-2,17-dioxa-4,9,15-triazanonadecane-5,10,14-tricarboxylate (6)

**5** (30 mg, 0.074 mmol. 1.0 eq.) and Fmoc-d-Glu-OtBu ( 31.7 mg, 0.075 mmol. 1.0 eq.) were dissolved in anhydrous DMF (1.5 mL). Then HBTU (31mg, 0.082 mmol, 1.1eq.) and HOBt (11 mg, 0.082 mmol, 1.1eq) were added and allowed to stir 5 minutes. DIPEA (52 μl, 0.29 mmol, 4 eq.) was added and reaction stirred overnight at room temperature. The reaction was then diluted with water and extracted with ethyl acetate (thrice). The combined organic fraction was then washed with 1N HCl (thrice), sat. sodium bicarbonate (thrice), brine (once) and dried over sodium sulfate. After concentration, the crude product was purified using flash chromatography (methanol/DCM/ 0.01% TEA-gradient from 1% to 5% methanol) to give **6** as an off-white solid (51 mg, 85%). ^1^H NMR (400 MHz, MeOD) δ 7.80 (d, *J* = 7.5 Hz, 2H), 7.73 – 7.64 (m, 2H), 7.39 (t, *J* = 7.5 Hz, 2H), 7.36 – 7.27 (m, 3H), 4.45 – 4.28 (m, 2H), 4.23 (t, *J* = 6.9 Hz, 2H), 4.08 (dd, *J* = 9.6, 4.8 Hz, 1H), 3.94 (dd, *J* = 9.0, 5.0 Hz, 1H), 3.00 (d, *J* = 6.7 Hz, 1H), 2.93 (d, *J* = 4.4 Hz, 1H), 2.57 – 2.38 (m, 0H), 2.32 (t, *J* = 7.6 Hz, 1H), 2.18 – 2.09 (m, 1H), 1.96 – 1.87 (m, 1H), 1.83 – 1.72 (m, 1H), 1.68 – 1.57 (m, 1H), 1.46 (s, 11H), 1.44 (s, 7H), 1.43 (s, 7H). ^13^C NMR (101 MHz, MeOD) δ 172.65, 142.40, 128.59, 127.97, 126.04, 120.73, 82.69, 82.61, 82.37, 67.79, 49.43, 49.21, 49.00, 48.79, 48.57, 48.36, 48.15, 32.08, 28.54, 28.08, 28.06, 23.06. LRMS (ESI) m/z: [M + H]^+^ Calcd for C_44_H_63_N_3_O_11_ 809.44; Found 810.44. NMR spectra shown in Figure S1F.

#### Tri-*tert*-butyl (6*R*,10*S*,15*R*)-2,2-dimethyl-4,12,17-trioxo-3-oxa-5,11,16-triazahenicos-20-yne-6,10,15-tricarboxylate (7)

**6** (57.8 mg, 0.071 mmol. 1.0 eq.) was dissolved in a solution of 20% piperidine in DMF (715 μL). The reaction stirred at room temperature for 4 hr. The reaction was diluted with water and extracted with ethyl acetate (thrice). The combined organic fraction was then washed with brine (once) and dried over sodium sulfate and concentrated and dried under vacuum. The crude product was then dissolved in anhydrous DMF (1.42 mL) and 4-pentynoic acid (11 mg, 0.106 mmol. 1.5 eq.) and EDC hydrochloride (34 mg, 0.177 mmol, 2.5 eq.) were added followed by DMAP (5.6 mg, 0.046 mmol, 0.65 eq.). The reaction stirred at room temperature overnight. The reaction was then diluted with water and extracted with ethyl acetate (thrice). The combined organic fraction was then washed with 1N HCl (thrice), sat. sodium bicarbonate (thrice), brine (once) and dried over sodium sulfate. After concentration **7** a was obtained as an off-white solid (32.7 mg, 34.4%, over two steps). LRMS (ESI) m/z: [M + H]^+^ Calcd for C_34_H_57_N_3_O_10_ 667. 40; found 668.40. Per literature precedent, further characterization is not performed [27].

#### (6*R*,10*S*,15*R*)-2,2-dimethyl-4,12,17-trioxo-5,11,16-triazahenicos-20-yne-6,10,15-tricarboxylate (iE-DAP-Alk) (8)

**7** (5 mg, 0.007 mmol, 1.0 eq.) was dissolved in 20% TFA in DCM (2.4mL) and the reaction stirred at room temperature for 6h. The reaction was then condensed, and the crude residue was triturated in cold diethyl ether. We note that the reaction must be carefully monitored and checked by NMR to assure that all 4 tert-butyl groups were removed. The crude product was precipitated, centrifuged, and purified with semi-prep C18 HPLC to give 9 as a white solid (1.0 mg, 30%). ^1^H NMR (600 MHz, D_2_O) δ 4.41 (dd, *J* = 8.8, 5.1 Hz, 1H), 4.36 (dd, *J* = 9.1, 5.3 Hz, 1H), 3.89 (t, *J* = 6.3 Hz, 1H), 2.56 – 2.50 (m, 2H), 2.45 (m, 2H), 2.40 – 2.37 (m, 1H), 2.24 – 2.16 (m, 1H), 2.08 – 2.02 (m, 2H), 2.01 – 1.87 (m, 3H), 1.80 (m, 2H), 1.58 – 1.46 (m, 2H).; HRMS (ESI-Pos) m/z: [M + H]^+^ Calcd for C_17_H_26_N_3_O_8_ 400.1714; Found 400.1744, see Figure 2 for proton NMR and high resolution Mass Spectra..

**Figure 2.**
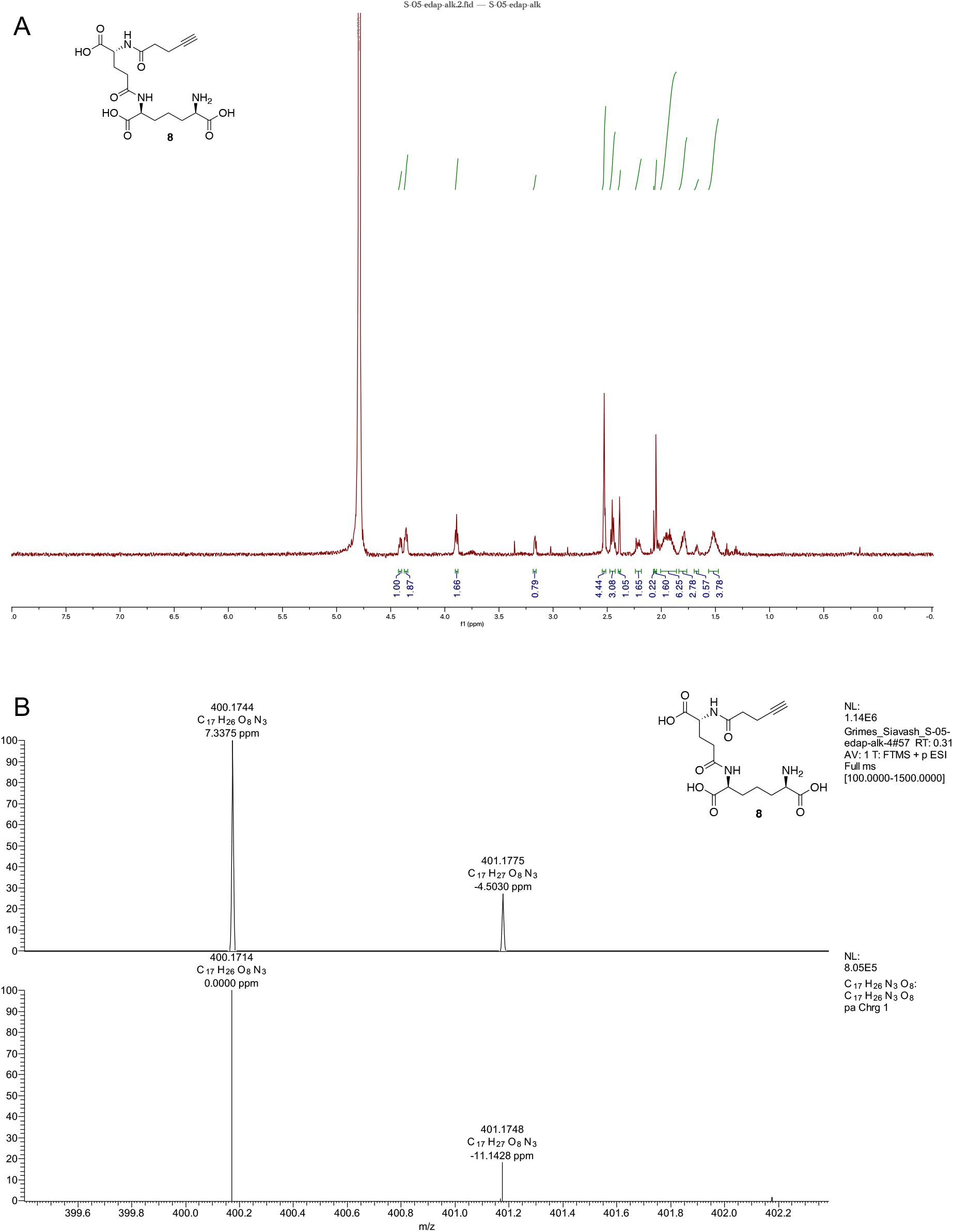
Spectral Data for iE-DAP-Alk. **(A)** 1H NMR **(B)** High Resolution Mass Spec Data Top: Experimental data Bottom: Simulated data made through Xcaliber.

### Synthesis of MDP-Alk

Synthesis of MDP-Alk follows an established synthesis of 2-amino-MDP [35] and instillation of the alkyne probe as the last step [27].

As shown in Figure 3, **(*R*)-5-amino-4-((*S*)-2-((*R*)-2-(((3*R*,4*R*,5*S*,6*R*)-2,5-dihydroxy-6-(hydroxymethyl)-3-(pent-4-ynamido)tetrahydro-2*H*-pyran-4 yl) oxy) propanamido) propanamido)-5-oxopentanoic acid (11):** 9 [36] (0.2136 g, 0.5142 mmol) was dissolved in anhydrous MeOH (7.3 mL) and Na_2_CO_3_ (0.2736 g, 2.5814 mmol) and **10** [37] (0.5016 g, 2.5701 mmol) was added in 5 portions over 1 h. Additional **10** (0.1009 g, 0.5170 mmol) was added at 24 h since the reaction was incomplete as determined by LC/MS. Once complete, the reaction was concentrated to dryness and then dissolved in water and IRA H+ resin was added. The filtrate was then collected and concentrated. The off-white solid was purified on semi-preparative HPLC The appropriate fractions were combined and lyophilized to give a white solid (0.1386 mg, 51% yield). ^1^H NMR (600 MHz, D_2_O): (Anomers 1.00α: 0.50β) δ 5.19 (d, *J* = 3.5 Hz, 1H), 4.71 (d, *J* = 8.4 Hz, 1H), 4.40 (dd, *J* = 9.7, 4.8 Hz, 1H), 4.34 (q, *J* = 6.8 Hz, 1H), 4.31 – 4.26 (m, 2H), 4.02 (dd, *J* = 10.5, 3.5 Hz, 1H), 3.97 – 3.64 (m, 5H), 3.63 – 3.53 (m, 3H), 3.53 – 3.47 (m, 1H), 2.53 – 2.42 (m, 6H), 2.39 – 2.34 (m, 1H), 2.26 – 2.18 (m, 1H), 2.08 – 1.89 (m, 1H), 1.49 – 1.43 (m, 3H), 1.41 – 1.35 (m, 3H);^13^C NMR (151 MHz, D_2_O) δ 176.96, 175.91, 175.70, 175.15, 174.65, 94.90, 91.02, 83.27, 81.77, 78.95, 77.51, 77.30, 75.68, 71.50, 70.17, 70.11, 69.11, 68.80, 60.76, 60.56, 56.20, 53.64, 52.60, 49.81, 34.84, 34.49, 30.06, 26.13, 18.40, 18.30, 16.60, 14.41, 14.33; HRMS (ESI-Pos) m/z: [M + H]^+^ Calcd for C_22_H_34_N_4_O_11_ 531.2297; Found 531.2299, see Figure 4 for NMR and high resolution Mass Spectra.

**Figure 3.**
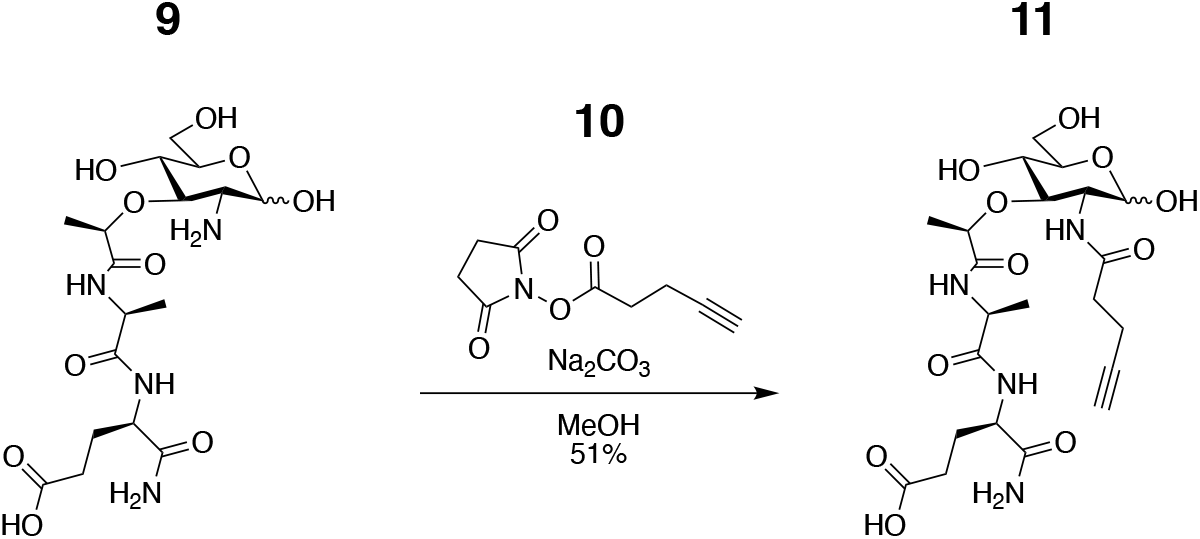
Schematic overview of the chemical synthesis of MDP-alkyne from the readily accessible 2-amino-MDP.

**Figure 4.**
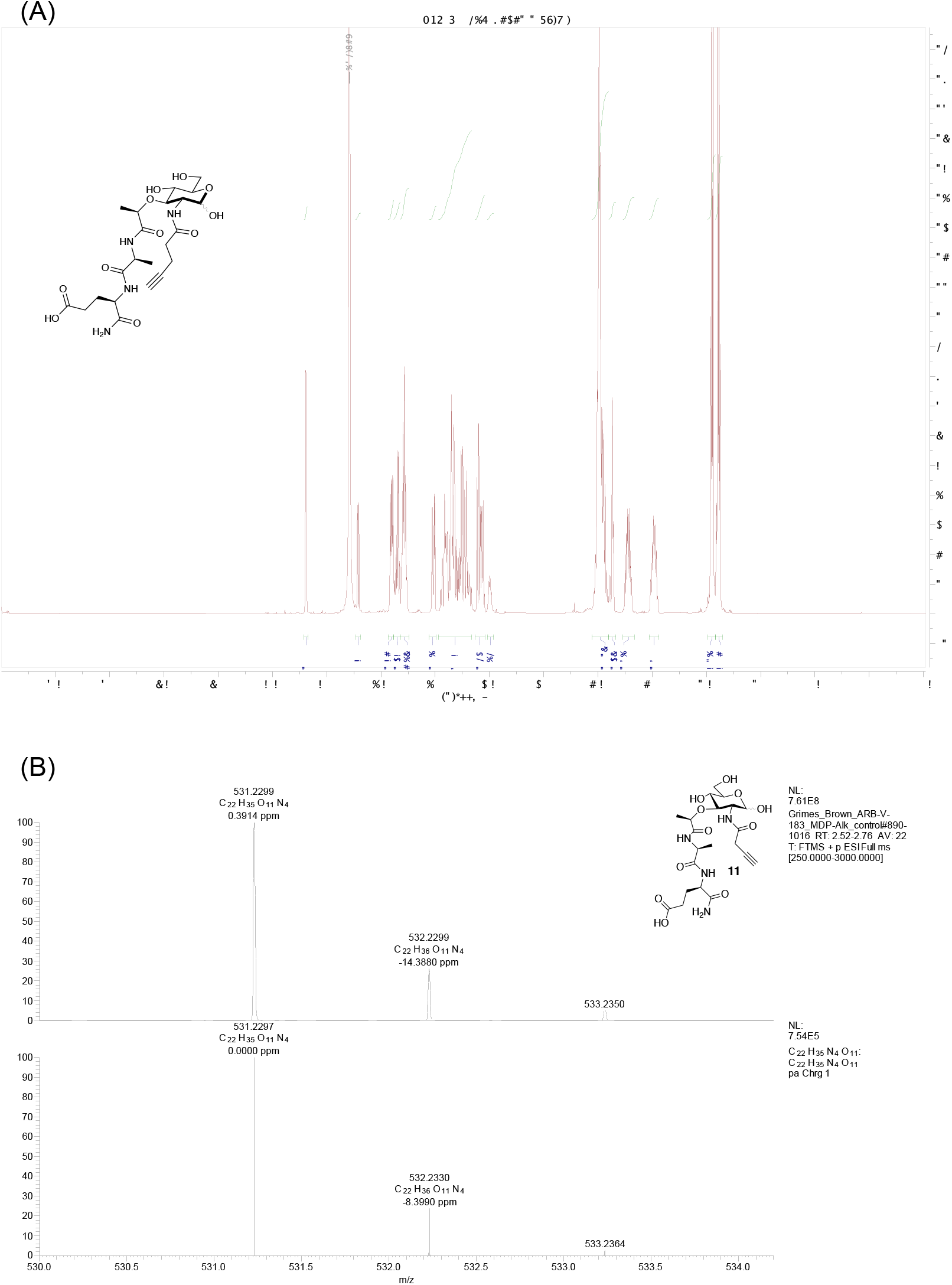
Spectral Analysis of MDP-Alk. **(A)** ^1^H NMR **(B)** High-Resolution Mass Spec Data Top: experimental data Bottom: Simulated data made through Xcaliber.

### Cell culture

HCT116 cells were isolated from a male human colorectal carcinoma and originally acquired from American Type Culture Collection (ATCC, CCL-247). HCT116 cells were maintained in DMEM (Corning CellGro) supplemented with 10% heat-inactivated FBS at 37°C, and 5% CO2.

### Dual-luciferase reporter assay

20,000 HCT-116 cells were incubated in 96-well plates and transfected with 50 ng NF-κB luciferase, 5 ng of pRL-TK (Promega), and 5 ng of pEF-Slc46 expression plasmid using GeneJuice (Millipore) for 24 h. In all cases, cells were stimulated by adding indicated agonist directly to the media for 6 or 24 h and subject to dual-luciferase assay. The TK-renilla was used to normalize the values from firefly luciferase. The values from unstimulated controls were subtracted from those for muropeptide-stimulated conditions to display the signal over the background. Each transfection was similarly analyzed in two independently transfected wells in a single plate, and all experiments were repeated with at least four independent assays [38].

### Mouse bone marrow-derived macrophages (BMDM)

Mouse BMDM were obtained from seven to twelve-week-old female mice maintained under pathogen-free conditions at the University of Massachusetts Medical School animal facilities, from C57BL/6 (WT), *Nod1^-/-^*, and *Nod2^-/-^* mice. Animals were sacrificed, femurs and tibia were dissected, and the bone marrow was flushed with PBS using a 30G needle connected to a 10 mL syringe. The cells were cultivated in RPMI (Corning) supplemented with 30% L929 cell-conditioned medium and 20% FBS (4×10^6^ cells/10 cm plate) at 37°C and 5% CO2. Cultures were re-fed on day 3 with the same media and maintained in RPMI supplemented with 5% L929 cell-conditioned medium and 10% FBS. BMDMs were used in the experiments on days 7–10 [39].

### BMDM stimulation protocol

BMDMs were primed with murine IFNγ (5 ng/ml, R&D systems) for 16 h, followed by the medium being replaced with fresh IFNγ (5 ng/ml) containing media for 2 h before treatment with indicated stimulations: MDP (100 μM, Invivogen), iE-DAP (100 μM, Invivogen) and ultra-pure Lipopolysaccharide (LPS) from *Escherichia coli* 0111:B4 (50ng/ml, Invivogen). After 24h stimulation media were processed for ELISA (R&D systems) as per manufacturer protocol [10].

### Microscopy

10^4^ cells (HCT116 or BMDM) were seeded on a glass coverslip in 24 well plates overnight. BMDMs were challenged with iE-DAP-Alk (30μM) or MDP-Alk (30μM) at 37 °C for 6h, cells washed twice with 1xPBS to remove access of click-muropeptides and fixed with 4% paraformaldehyde in PBS at RT for 20 min. Cells were permeabilized with 0.1% Tween-20 in PBS for 10 min at RT and blocked with 1% BSA in PBS. These permeabilized cells were then incubated in click-reaction conditions (250μM CuSO4, 35μM BTAA, 60μM sodium ascorbate) with 2.5μM CalFluor 488 Azide at RT for 30 mins. Cells were then washed with DPBS thrice and mounted on slides, with DAPI containing mounting media. Slides were imaged with a Leica SP8 confocal microscope.

## Results

### Validation of NF-κB stimulating activity

First, we checked the potency of iE-DAP-Alk and MDP-Alk using an established NF-κB luciferase assay [38]. We expressed by transient transfection human *Slc46A2* in HCT116 colon carcinoma cells, which are known to display enhanced responses to NOD1/2 ligands when these SLC46s are expressed [38]. These cells were challenged with multiple doses of native iE-DAP, iE-DAP-Alk, native MDP, or MDP-Alk for 6hrs and 24hrs (Figure 5). iE-DAP and iE-DAP-Alk showed robust and similar induction of NF-κB reporter activity. MDP and MDP-Alk also showed an almost identical does responses in NF-κB reporter activity (Fig.5). Results from these experiments confirm that iE-DAP-Alk and MDP-Alk have similar potency as the native molecules in triggering the NOD1/NOD2 NF-kB pathway.

**Figure 5.**
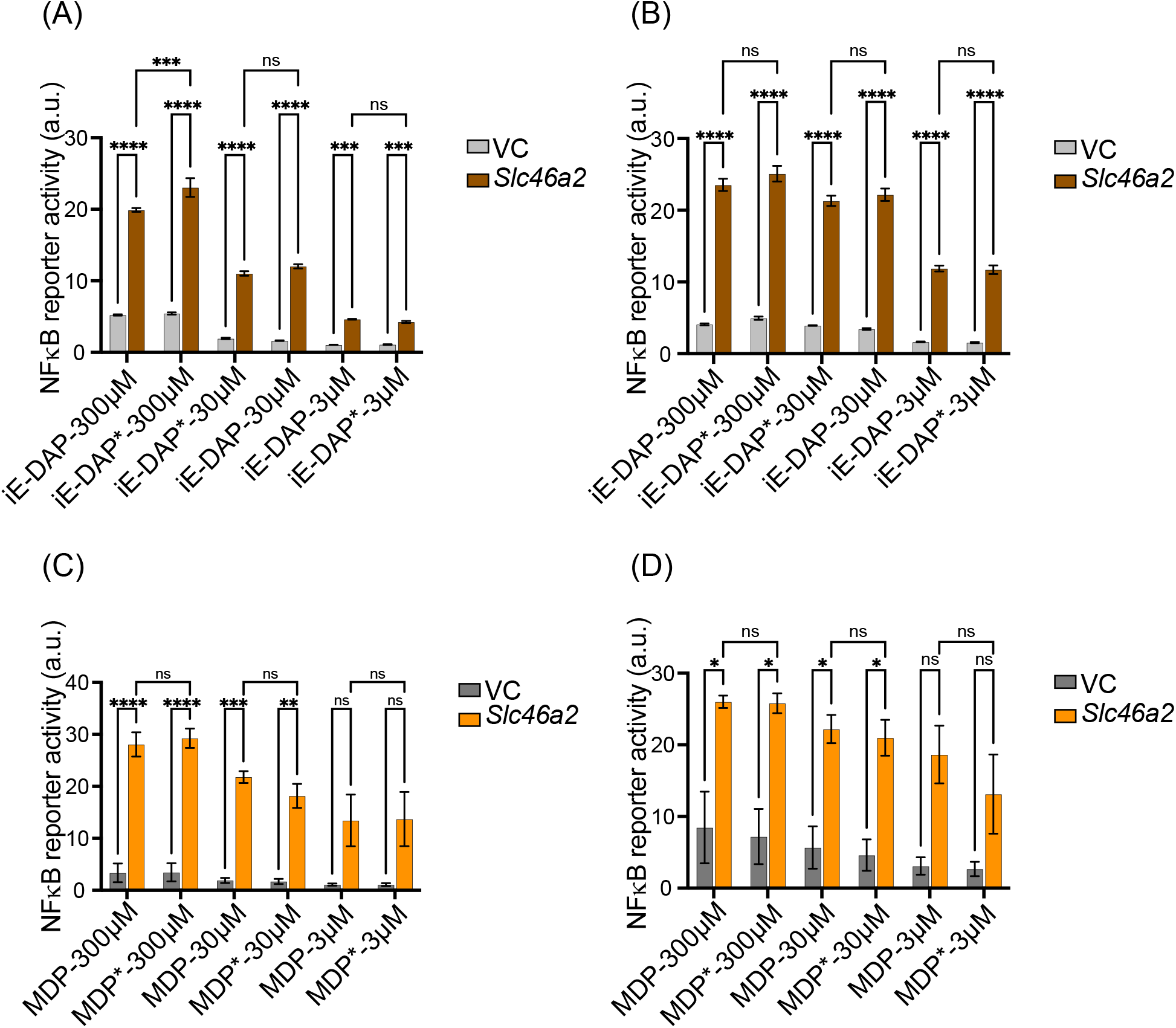
Alkyne-modified iE-DAP and MDP showed similar potency in cell line reporter assays. NF-κB luciferase assays in HCT116 cells transiently expressing human *Slc46a2* or empty vector (VC) showed that multiple doses of native iE-DAP/iE-DAP-Alk (iE-DAP*) and native MDP/MPD-Alk (MDP*) triggered indistinguishable NF-κB reporter activity at 6 or 24 h, A&C or B&D respectively. Values from unstimulated controls were subtracted from the values for muropeptide-stimulated conditions, to display the signal over the background. N = 4, two-way ANOVA with Tukey’s multiple comparisons test to determine significance. **** P < 0.0001; *** P < 0.001; ** P < 0.01; * P < 0.05; ns, not significant.

### BMDM cytokine stimulation assay

iE-DAP-Alk and MDP-Alk potency were further tested with IFNγ-primed bone marrow-derived macrophages (γ-BMDM) by measuring interleukin-6 (IL-6) and tumor necrosis factor (TNF) production [11]. BMDM from WT, *Nod1^-/-^* or *Nod2^-/-^* were challenged with native and iE-DAP-Alk and MDP-Alk (100 μM). Both native and alkyne-modified molecules similarly stimulated IL-6 and TNF production in WT γ-BMDM culture (Figure 6). Negligible cytokine production was observed in *Nod1* or *Nod2* deficient γ-BMDM challenged with iE-DAP or MDP, respectively (See Figure 6). These results demonstrate that the modified molecules have similar activity to the native NOD1/2 agonists. LPS was used a control cytokine inducing stimuli.

**Figure 6.**
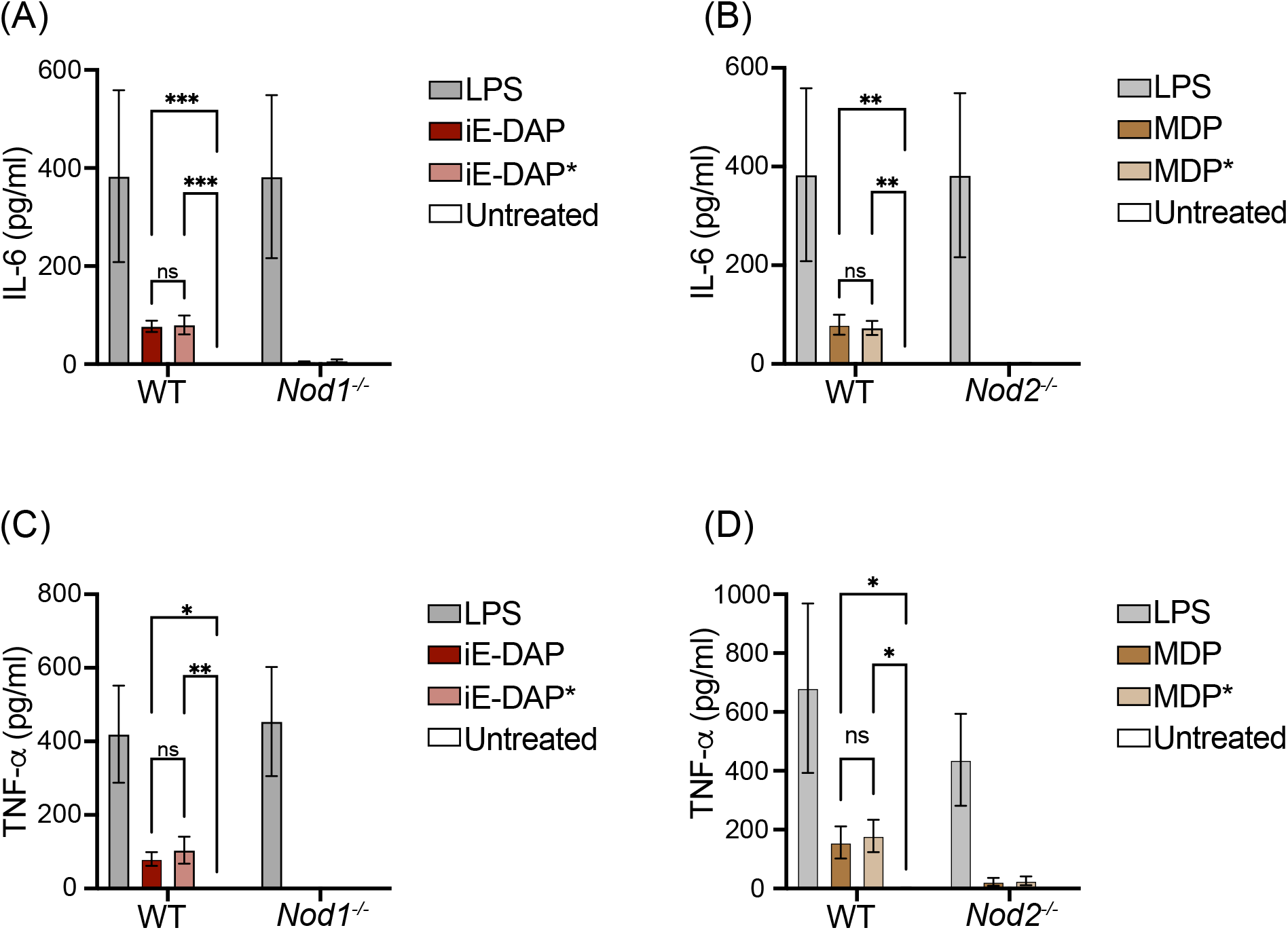
Alkyne -and native iE-DAP and MDP stimulate similar levels of IL-6 and TNF-α in macrophages. **(A** and **B)** IFNγ-primed BMDMs with indicated genotypes were challenged with different agonists as labeled at 100 μM, and IL-6 was measured by ELISA. (C and D) TNFα was measured by ELISA after being challenged with different agonists as in Figures A and B. N= 3, two-way ANOVA with Tukey’s multiple comparisons test to determine significance. **** P < 0.0001; *** P < 0.001; ** P < 0.01; * P < 0.05; ns, not significant.

### Internalization of modified iE-DAP and MDP

iE-DAP and MDP are recognized by intracellular receptors NOD1 and NOD2 respectively. Therefore, their delivery to the cytosol is critical for downstream signaling [10, 11]. To confirm the internalization of modified molecules into the cell, IFNγ-primed BMDMs were challenged with iE-DAP-Alk and MDP-Alk and visualized using click chemistry, with azide-linked Calflour-488 (AZDye™-488). Both iE-DAP-Alk and MDP-Alk were observed in the cytosolic compartment of WT as well as *Nod1*- and *Nod2*-deficient BMDMs after a 6h challenge. Results from these experiments confirm the delivery of modified iE-DAP and MDP into the intracellular compartments of BMDMs. Moreover, it also shows that intracellular delivery of iE-DAP or MDP is not dependent on *Nod1* or *Nod2* (Figure 7A). We have further confirmed these results in HCT116 cells transiently expressing mouse *Slc46A2* or *Slc46A3*. It was previously shown that HCT116 cells respond weakly to muropeptides at baseline, but *Slc46A2* or *Slc46A3* expression strongly enhanced responses to these NOD1/2 agonists ([38], and Figure 2). HCT116 cells expressing *Slc46a2* or *Slc46a3* imported iE-DAP-Alk and MDP-Alk, respectively, using azide-linked Calflour-488 (AZDye™-488) and confocal microscopy for detection, whereas empty vector control transfected cells did not show any sign of internalization of these modified muropeptides (Figure 7B). Results from these experiments confirm that alkyne modification did not affect the functionality or activity of iE-DAP and MDP, while these molecules are useful for cell biological analysis of muropeptide trafficking.

**Figure 7.**
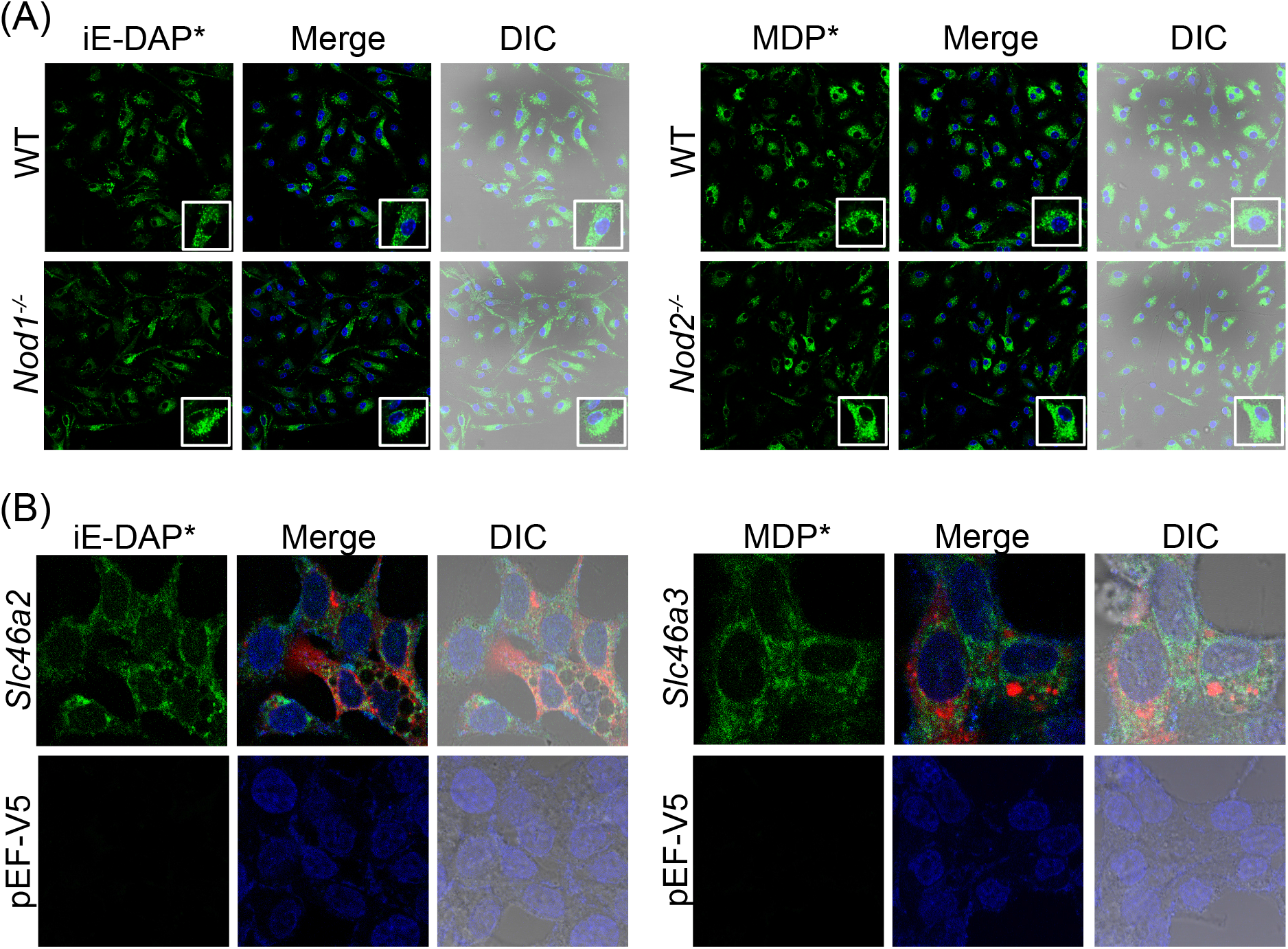
Internalization of iE-DAP-Alk and MDP-Alk in BMDMs and HCT116 cells. **(A)** Using click-chemistry with azide-Calflour (green), the intracellular distribution of iE-DAP-Alk or MDP-Alk, independent of *Nod1* or *Nod2*, was observed by confocal fluorescent microscopy in bone marrow-derived macrophages. Nuclei stained with Hoechst (blue) **(B)** HCT116 cells transiently expressing mouse *Slc46a2* or *Slc46a3* similarly showed intracellular iE-DAP-Alk or MDP-Alk uptake (green), whereas empty vector-transfected cells failed to import these NOD1/2 agonists. Red stain is V5 tagged Slc46a2 or Slc46A3. Nuclei stained with Hoechst (blue). Images in panels are representative of three independent experiments.

## Discussion and Conclusion

The microbiota is an essential component of animal anatomy and physiology and plays a critical role in human health [40]. Bacteria are the most abundant microorganism in the microbiota and play a significant role in maintaining homeostasis with innate and adaptive immune functions [41, 42]. Immune receptors NOD1 and NOD2 play a major function in detecting bacteria or bacterial components, especially at barrier layers [43]. However, much still needs to be learned about how these receptors specifically recognize muropeptides and how these muropeptides gain access to these cytosolic receptors. iE-DAP and MDP have been identified as minimal agonists for NOD1 and NOD2, respectively. Here, we report the synthesis and validation click reaction-supporting alkyne-linked iE-DAP and MDP. While the synthetic process can be extensive and time-consuming, synthesizing the fragments in-house allowed us to control the stereochemistry of the molecule [44–46]. For example, in the synthesis of *m*-DAP we are able to selectively attain one enantiomer whereas commercially available iE-DAP is a mixture of both enantiomers. In the future, these “click-able” probes will be further modified, beyond the alkyne handle, to allow us to functionalize for further biological probing.

Using an established NF-κB luciferase reporter assay, where expression of SLC46s in HCT116 cells supports robust NF-kB activity upon iE-DAP or MDP challenge [38], we have compared the potency of iE-DAP-Alk and MDP-Alk with the native molecules. HCT116 cells express both NOD1 and NOD2, making them a suitable option to test both iE-DAP and MDP [47, 48]. These assays showed very similar activities for the alkyne-modified muropeptides, compared to their native counterparts, at multiple doses and two different time points.

Further, we assessed the innate immune stimulatory activity of iE-DAP-Alk and MDP-Alk in IFNγ-primed mouse BMDMs and again found similar cytokine promoting activity compared to their native counterparts. Modified molecules elicited the same level of IL-6 and TNF in BMDMs as the native NOD1/2 agonists. The same system (γBMDMs) as well as *Slc46a2/a3* transfected HCT116 cells were utilized to demonstrate the internalization of modified iE-DAP and MDP, taking advantage of the click handles. Click-chemistry is particularly useful to visualize the trafficking of small molecules like iE-DAP or MDP in cellular compartments, using an azide-linked fluor substrate in the presence of the copper catalyst. The substrates for the click azide reaction are very specific to the reactant and this azide-dye reaction is only catalyzed in the presence of copper. No free alkyne or azide group is found in biomolecules, which makes the click reactions very precise [49, 50]. Overall, the results shown here demonstrate that alkyne-modified iE-DAP and MDP are similarly active as native muropeptides in eliciting NF-kB activity and or cytokine production. Moreover, these alkyne-modified muropeptides are valuable for visualizing their localization and trafficking within intracellular compartments of mouse BMDM and human cell lines.

## Notes

### Competing Interest Statement

A provisional patent on targeting SLC46s to inhibit inflammation in psoriasis and other auto-inflammatory diseases as been filed by some of the authors.

