## Supplemental Figure for "Synthesis and validation of click-modified of NOD1/2 agonists"

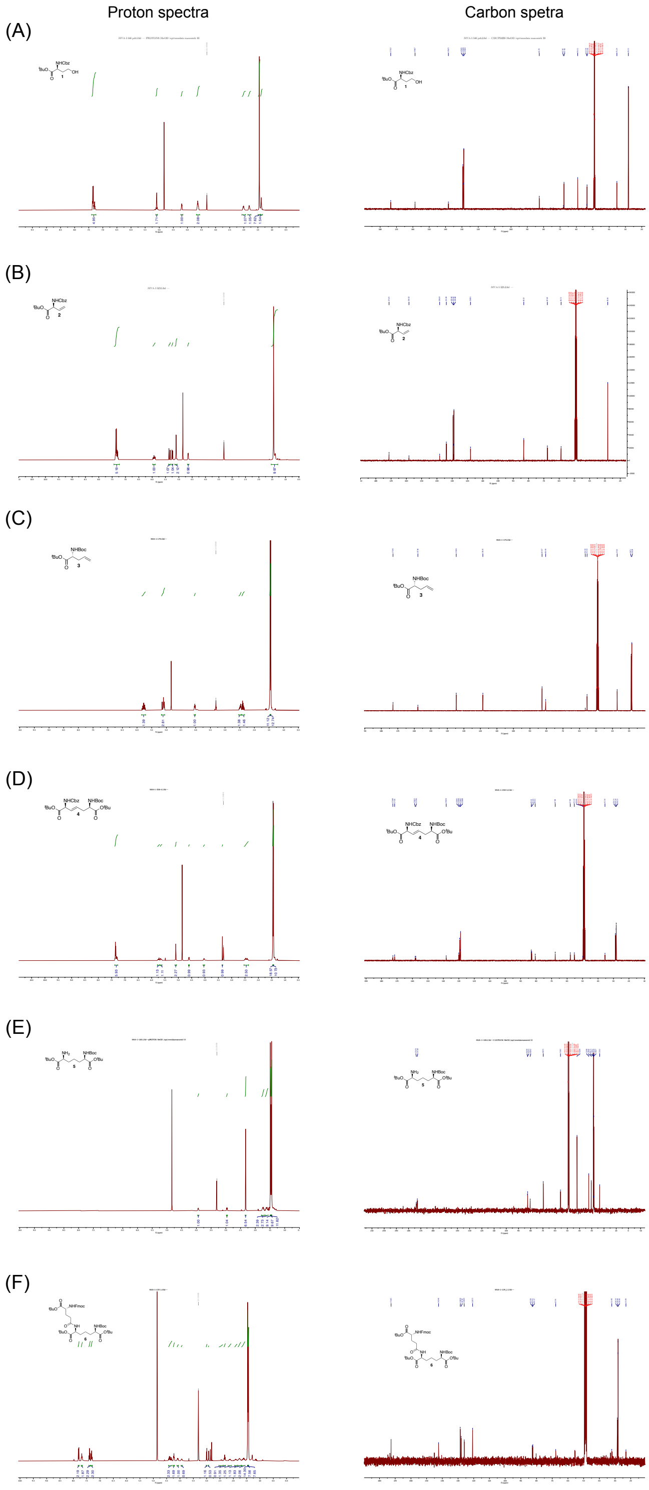


**Figure S1. Proton and carbon NMR spectra obtained from the iE-DAP-Alk synthesis reported in this manuscript**. As the synthesis of this molecule is demanding, these spectra of intermediates to the final compounds are provided as a resource for those embarking in the production of these molecules. The details provided here will assist in future production of PGN derivatives.
